# Toxicogenomic analysis of the carcinogenic potential of plastic additives

**DOI:** 10.1101/2023.06.30.547246

**Authors:** Sophia Vincoff, Beatrice Schleupner, Jasmine Santos, Margaret Morrison, Newland Zhang, Meagan M. Dunphy-Daly, William C. Eward, Andrew J. Armstrong, Zoie Diana, Jason A. Somarelli

## Abstract

Plastics are the most prevalent human-made substance in the world and are ubiquitous throughout all ecosystems. Microscopic plastic particles are routinely ingested and inhaled by humans and other organisms. Despite the frequency of plastic exposures, little is known about their health consequences. Of particular concern are plastic additives -chemical compounds that are mixed into plastics to improve functionality or are unintentionally acquired during plastic production and use. Additives are loosely bound to the plastic polymer and may be released during plastic exposures. These compounds may pose health concerns, such as cancer, but little is known about the potential impact of these chemicals on health. To better understand the health effects of plastic additives, we performed an integrated toxicogenomic analysis on 2,712 additives, focusing on cancer as a well-studied toxicological endpoint. Screening these substances across three chemical databases revealed two key observations: 1) over 150 plastic additives have known carcinogenicity and 2) the majority (∼90%) of plastic additives lack data on carcinogenic endpoints. Analyses of additive usage patterns pinpointed specific polymers, functions, and products in which carcinogenic additives reside. Based on published chemical-gene interactions, both carcinogenic additives and additives with unknown carcinogenicity impacted similar biological pathways. The predominant pathways involved DNA damage, apoptosis, immune response, viral diseases, and cancer. This study underscores the urgent need for systematic and comprehensive carcinogenicity assessment of plastic additives and regulatory responses to mitigate the potential health risks of plastic exposure.

## Introduction

Plastics, over the last half-century, have established a worldwide presence in nearly all societies and are widely detectable as pollutants in the environment. Demand for plastics has skyrocketed since the 1950s due to their inexpensive, strong, durable, and lightweight properties ^1^. Between 1950 and 2015, 8.3 billion metric tons of plastic products were created worldwide, 75% of which became waste ^2^. Global plastic production is expected to double by 2040 ^3–5^. Despite benefits to societal health and safety, as well as a global market value of USD 569.9 billion in 2019 ^4,6^, there is growing concern about plastic pollution and its impact on human and organismal health ^7–13^.

Humans regularly interact with plastics through food packaging, clothing, toiletries, household items, furniture, automotives, medical equipment, electronics, toys, and office supplies ^14^. All plastics experience weathering, leading to the release of microplastics (1μm-5mm) and nanoplastics (<1μm) ^15–17^. While initial human interactions with plastics are typically by choice, the ubiquitous persistence of plastics in the environment means that many subsequent exposures are involuntary. For instance, humans are routinely exposed to plastic particles through respiratory, oral, and dermal routes ^18–20^. As a result, plastic has been detected in human tissue and secretions, such as the lungs, colon, breast milk, and placentas ^21–24^. Given the widespread presence of plastics and microplastics in the environment and human bodies, there is an urgent need to determine the health impacts of plastics. At present, there is far more information regarding exposure to individual polymers or specific plastics than there is about the lifetime exposure to all plastics^25^.

The risks of plastic exposure cannot be assessed without first acknowledging that plastics are not pure substances, but rather complex mixtures of polymers along with dozens to thousands of chemical compounds broadly categorized as additives ^26,27^. Common additives used for performance enhancement include plasticizers, flame retardants, heat and light stabilizers, antioxidants, lubricants, pigments, antistatic agents, slip agents, biocides, and thermal stabilizers ^28^. Plastics also contain non-intentionally added substances from manufacturing, such as residual monomers, byproducts, and contaminants ^14^. During and after plastic usage, additional substances are adsorbed from the environment ^29^, such as polycyclic aromatic hydrocarbons or alkylphenols ^30^.

Whether intentionally incorporated or not, plastic additives have the potential to leach from plastics and contaminate soil, air, water, food, and human bodies ^31^. Additives can comprise a sizable mass fraction of a plastic polymer ^32^, such as plasticizers which are incorporated up to 70% weight by weight in some polymers ^31^. Plastic additives have been detected in biota and throughout the environment, including in the tissues of shellfish ^33^, fish ^34,35^, seabirds ^36^, and marine mammals ^37^, underscoring the need to elucidate the impacts of these chemicals on organismal health. For instance, plastic particles and their additives have been linked to cancer and carcinogenic processes ^18,28,29,38,39^; however, a comprehensive analysis of the carcinogenic potential of plastic additives has not been performed.

Previous studies have identified many commonly used plastic additives, including those often used in food-contact products as well as those that should be further studied for their potential impacts on organismal health ^28,31^. Other studies have begun to identify the additives used in particular sectors of the plastic industry (*e*.*g*., packaging), but thousands of additives remain uncharacterized ^14,28,31^. Thus, no comprehensive list of plastic additives exists, nor are there established standards for additive hazard classifications or regulations ^14^.

Here we investigate the carcinogenic risk of plastic additives and identify gaps in the current knowledge. To do this we curated a list of over 2,700 additives through a literature search of three databases. By querying three public chemical registries, we identified those additives with known and unknown carcinogenic potential. Using a toxicogenomics approach, we assessed the potential mechanisms of carcinogenicity and identified enriched pathways for all additives. The majority of our additives (∼90%) were unclassified as to their carcinogenicity in two major registries, either due to a lack of toxicological data or no public concern over the danger of the chemical. However, of the 229 unclassified additives with enough published gene expression data for analysis, a substantial portion (80.3%) induced genetic pathways related to cancer and cancer-like phenotypes. Together, these analyses demonstrate a dearth of public knowledge regarding plastic additive carcinogenicity and pinpoint the need for a comprehensive experimental framework to determine the toxicological effects of plastic additives.

## Results

This study resulted in a methodological workflow (**Fig. 1**) to compile a list of plastic additives, investigate the carcinogenicity classifications of the additives, determine known impacts on gene expression, predict additives’ interference with human biological pathways, and group additives according to their predicted pathway effects. An abbreviated parallel analysis was conducted on 281 reported polymer backbones, such as polyvinyl chloride and latex (**Table S2**). All collected data regarding additives can be found in Table S1.

**Figure 1:**
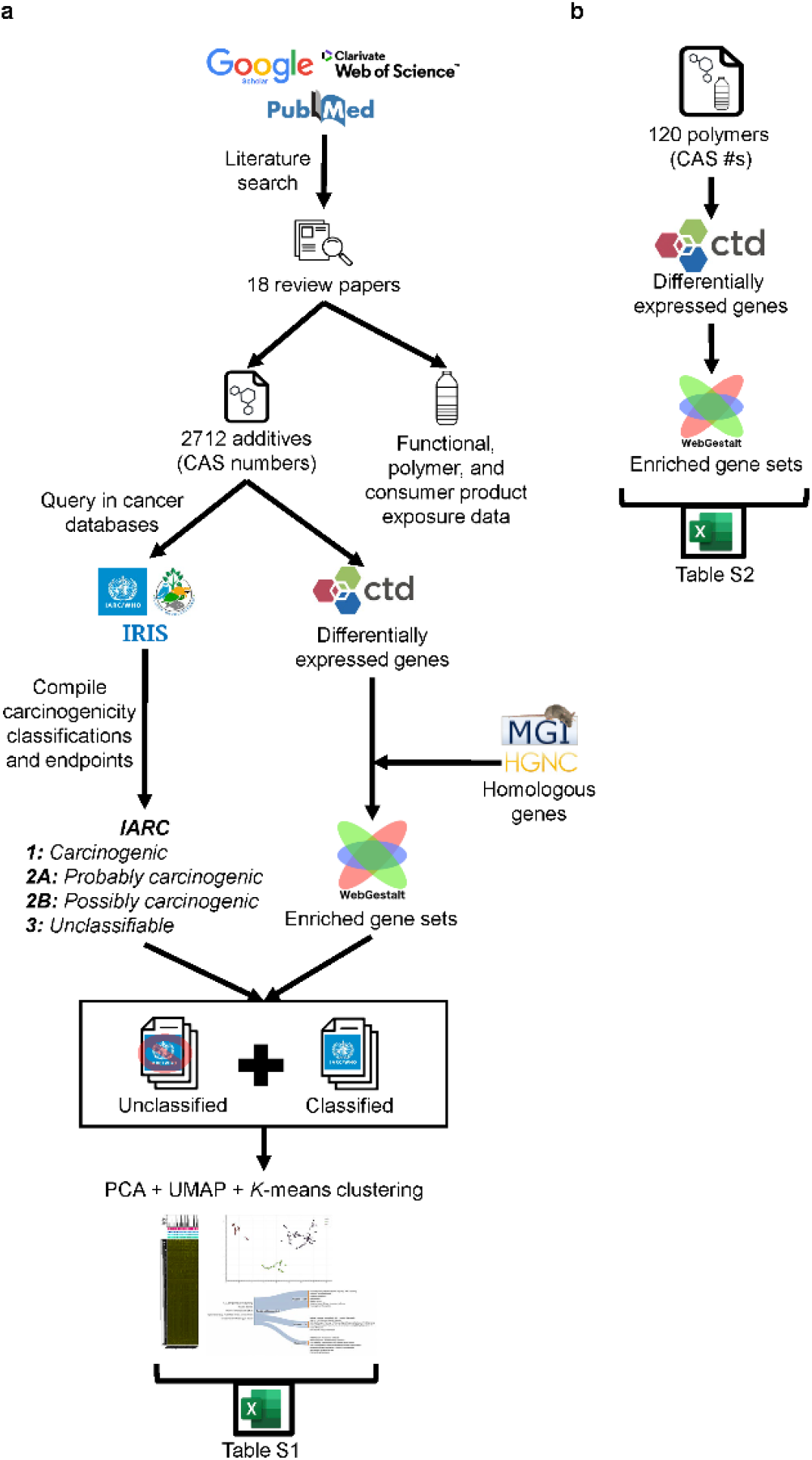
Resulting methodological workflow to analyze the carcinogenicity and gene expression patterns of **a**) plastic additives and **b**) polymers.

### Plastic additives include multiple known carcinogens and many with unknown cancer-causing potential

We first examined the presence and classifications of additives within the International Agency for Research on Cancer (IARC), which contained 1,101 chemicals at the time of our analysis (**Fig. 2a**). A total of 2,421 additives (89.27%) were absent from IARC (**Fig. 2a**). Among the 291 additives in the database, 12 (4.12%) had no classification, 112 (38.5%) had inadequate evidence for carcinogenicity and require more research (Group 3), 108 (37.1%) were possibly carcinogenic (Group 2B), 36 (12.4%) were probably carcinogenic (Group 2A), and 23 (7.9%) were carcinogenic (Group 1).

**Figure 2:**
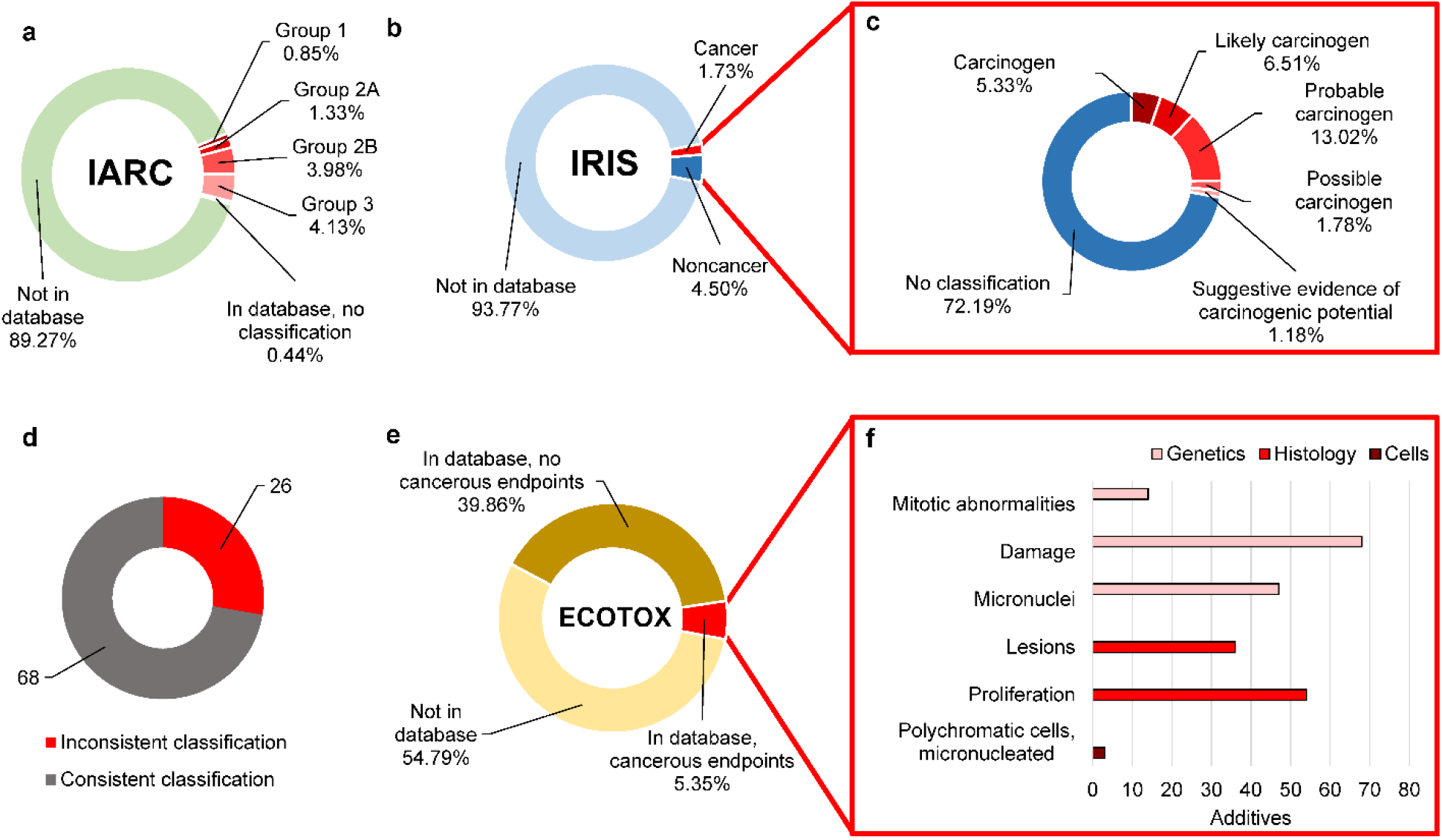
The majority of plastic additives are undocumented in toxicological databases. **a)** Classification of the 2,712 plastic additives in the IARC database (Group 1: carcinogenic, Group 2A: probably carcinogenic, Group 2B: possibly carcinogenic, Group 3: not classifiable as to its carcinogenicity in humans). **b)** Classification of plastic additives in the IRIS database (Cancer = evidence of carcinogenicity or known carcinogen according to IRIS. Noncancer = chemical in the IRIS database, but no evidence suggesting carcinogenicity). **c)** Detailed classifications within the ‘cancer’ and ‘noncancer’ categories in the IRIS database. **d)** Consistency of the classified additives in IRIS and IARC. **e)** Classification of plastic additives in the ECOTOX database. **f)** Cancerous endpoints among the 5.35% of plastic additives with data in the ECOTOX system.

Cross-referencing all 2,712 additives in the IRIS database revealed 2, 543 (93.77%) with no cancer-related data, 122 (4.50%) listed as noncancerous, and 47 (1.73%) listed as cancerous (**Fig. 2b**). The IRIS database contained 651 chemicals at the time of our analysis. Of the 169 additives present in IRIS, 9 (5.33%) were carcinogens, 11 (6.51%) were likely carcinogens, 22 (13.02%) were probable carcinogens, 3 (1.78%) were possible carcinogens, and 2 (1.18%) had suggestive evidence of carcinogenicity (**Fig. 2c**). Ninety-four additives were present in both the IARC and IRIS databases (**Table S1**), 26 of which had inconsistent cancer classifications between the two databases (**Fig. 2d**).

In the ECOTOX database, 1,486 (54.79%) were not present, 1,081 (39.68%) were present in the database without cancer-relevant endpoints, and 145 (5.35%) were present in the database with cancer-relevant endpoints (**Fig. 2e**). Each of those 145 additives contained one or more cancer-relevant endpoints. In total, 68 additives were associated with genetic damage endpoints, 54 with proliferation, 47 with micronuclei, 36 with lesions, 14 with mitotic abnormalities, and 3 with micronucleated polychromatic cells (**Fig. 2f**).

### Additive exposure data are sparse, particularly when carcinogenicity is uncertain

We next collected exposure information for all plastic additives with available data (2508, 94.28%) from 18 review papers (**Fig. 1)**. In our analysis, *classified* additives are those assigned to Group 1, 2A, 2B, or 3 in IARC, *unclassified* additives are those absent from or unassigned in IARC, and *carcinogenic* additives are the subset of classified additives in Group 1, 2A, or 2B.

Analysis of exposure data indicated 1,477 total additives (184 classified, 1,293 unclassified) associated with at least one polymer, 2,315 additives (248 classified, 2,067 unclassified) with at least one functional annotation, and 892 additives (104 classified, 788 unclassified) associated with at least one industrial or consumer product (**Fig. 3a**). In total, 546 additives (84 classified, 462 unclassified) have exposure data in all three categories (product, function, and polymer).

**Figure 3:**
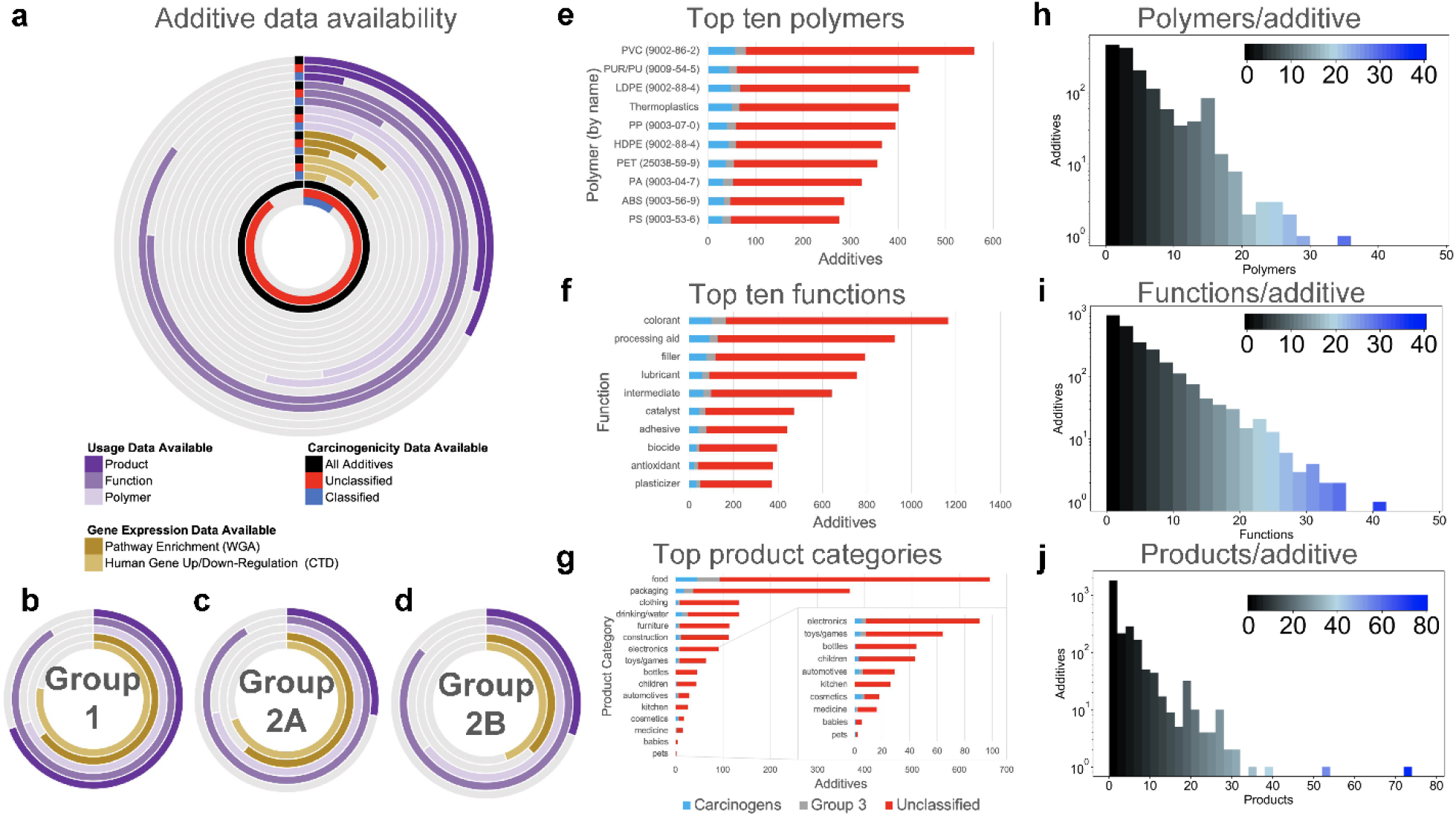
Data on usage, exposure, and health effects for 2,712 plastic additives. **a)** Product associations, polymer associations, effects on human gene expression, and possible disturbances to human gene networks for plastic additives. Dark-colored lines indicate the proportion of additives (out of 2,712) for which information is available. Rings marked or colored black represent all additives; red represents unclassified additives; blue represents classified additives. **b-d)** Knowledge of additive carcinogenicity is associated with more knowledge of biological properties (**Table S5**). **e-g)** The ten polymers, functions, and product categories with the most additive associations. Polymer associations were determined by Reference Name (**Table S2**). ***\*****LDPE and HDPE share the same CAS number*. **h-j)** Distributions of polymers/additive, functions/additive, and products/additive. Additives with no associations to polymers, functions, or products are not included in the histogram.

Fewer than one-third of all carcinogenic additives are linked to industrial or consumer products (**Fig. 3b-d**). Nine of the top ten polymers by additive association have traceable Chemical Abstracts Service Registry Numbers (CASRNs, or CAS numbers), and each is connected to hundreds of additives (**Fig. 3e**). Nearly 400 additives are listed as components of “thermoplastics,” which encompass all plastics that become moldable at high temperatures and solidify upon cooling, including acrylic, polypropylene (PP), and polystyrene (PS). The top ten functions out of 167 unique function strings are reported in **Fig. 3f**, with the top four (colorant, processing aid, filler, lubricant) mapped to over 600 additives each. Sixteen product categories of interest were extracted from the product data by querying our database with specific search strings (**Table S3**). Food, packaging, and clothing-related products are associated with the most additives; medicine, babies, and pets are associated with the fewest (**Fig. 3g**).

Regarding the polymer data, we found that each additive is associated with 2.4 + 3.9 polymers on average **(Fig. 3h)**, and >200 additives are associated with over 10 polymers each. Diethylhexl phthalate, a type 2B carcinogen (CAS = 117-81-7) has the maximum polymer associations (34) and 18 documented functions (**Table S1**). This additive was linked to diverse products including food packaging, plastic bags, medical equipment (*e*.*g*., syringes, dialysis equipment, catheters, intravenous tubing, blood/dialysis bags, gaskets, implants, gloves), baby products (*e*.*g*., pacifiers), plastic toys (*e*.*g*., soft squeeze toys, balls, light sticks), bathroom products (*e*.*g*., shower curtains, sanitary products), leisure products (*e*.*g*., colored fishing floats, sports equipment), clothing (*e*.*g*., raincoats), furniture (*e*.*g*., floor tiles, furniture upholstery, car seats, tablecloths, flooring, wall coverings, wood coatings), and articles intended for pets.

The majority of additives are associated with up to five functions and products (**Fig. i-j**), but several additives have dozens of matches in at least one exposure category. Formaldehyde, a Type 1 carcinogen (CAS = 50-00-0), is the most function-heavy, with 40 documented functions. This chemical also features 17 polymer associations and nine product associations, including food contact products, manufacturing container metals, and car seat stuffing. Similarly, butylated hydroxytulouene (CAS = 128-37-0; a class 3 chemical in IARC), and bisphenol A (80-05-7; a chemical unclassified in IARC) have very high numbers of both function and polymer associations (**Table S1)**.

### Plastic additives impact diverse gene expression pathways

We used the up- and downregulated genes associated with all plastic additives in the Comparative Toxicogenomics Database (CTD) (18782832) as inputs for over-representation analyses in WebGestalt (WGA) (31114916). The most commonly upregulated genes by plastic additives include the tumor suppressor TP53, the pro–inflammatory cytokines C-X-C Motif Chemokine Ligand 8 (CXCL8, IL-8) and CXCL6 (IL-6), genes responsible for detoxification and metabolism of toxins, such as CYP1A1, and the cell cycle regulator, CDKN1A. The genes downregulated by the greatest number of additives included the apoptosis regulators, BCL2, BCL2L1, and BAX, and the cell adhesion molecule and epithelial lineage marker, E-cadherin (CDH1). Whether additives are classified or unclassified in regard to carcinogenicity, the reported effects on gene expression are similar (**Table S4)**.

At the pathway level, carcinogenic and unclassified additives have similar impacts. Pathways altered by both carcinogens and unclassified additives include apoptosis, pathways in cancer, and signaling by interleukins. However, unclassified additives, but not carcinogens, alter cytokine signaling in immune system, lung fibrosis, and the AGE-RAGE signaling pathway in diabetic complications (**Table S4**).

Of the 2,712 additives, only 428 (15.78%, 139 classified, 289 unclassified) modulated human gene expression according to the CTD, and 349 additives (12.87%, 120 classified, 229 unclassified) contained enough gene interactions for over-representation analysis (**Fig. 3a**). As IARC predictions intensified, from unclassified to Group 1, the number of papers documenting chemical-gene interactions increased (**Fig. 3a-d, Table S5**). Group 1 carcinogens were found to have significantly more gene interaction data (p < 0.05) than any other group (**Table S5)**.

### Based on pathway enrichment ratios, classified and unclassified additives cluster into three unique groups

We next used *K*-means and hierarchical clustering to visualize the relationships between additives at the pathway level (**Fig. 4**). Pathway over-representation enrichment ratios (ERs) were used as input for the clustering. Clustering on all additives and all ERs produced silhouette scores indicating that *k=3* clusters were optimal (**Table S5**). This clustering was largely unchanged when we analyzed subsets of data by similar pathway names (*e*.*g*., containing substrings of cancer keywords [‘cancer’, ‘carcin’, ‘metasta’, ‘tumor’] (**Fig. 4b)** or oxidative stress keywords (**Fig. S2**)), indicating that the clusters are well-separated. Notably, although each of the clusters are of different sizes, all clusters contain similarly-proportioned mixtures of carcinogens (22-24%) and unclassified additives (65-69%) (**Fig. 4c-e**), suggesting that carcinogenic and unclassified additives impact gene expression in similar ways (Cluster 1: 143 unclassified, 49 carcinogenic; Cluster 2: 51 unclassified,17 carcinogenic; Cluster 3: 35 unclassified, 12 carcinogenic). Additives within each cluster also exhibit diverse exposure data. A mixture of Group 1, 2A, 2B, 3, and unclassified additives are present in PVC (the most common polymer), used as colorant (the most common function), and/or found in food products (the most common product category), but the specific proportions vary by cluster (**Fig. 4c-e**).

**Figure 4:**
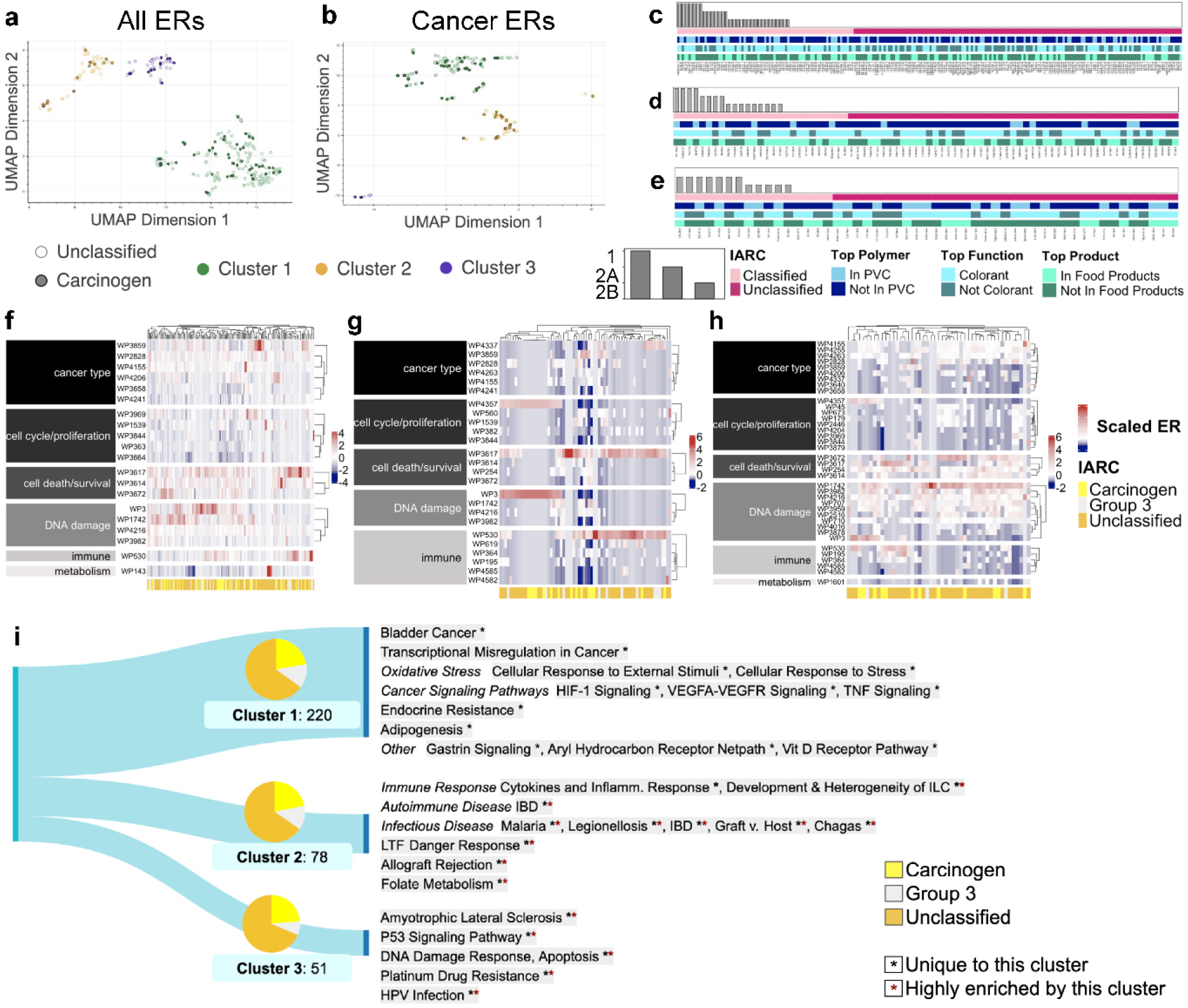
Carcinogens (probable, possible, or confirmed) and unclassified additives share similarities in their impacts on gene expression. When additives are clustered on their ERs for KEGG, Reactome, Wikipathways, and PANTHER gene sets, they form three distinct groups containing near-identical distributions of carcinogens, Group 3 additives (inadequate evidence for carcinogenicity in humans), and unclassified additives. **a)** Three clusters (silhouette score = 0.86) encompass all additives, based on ERs for all 2,246 pathways. **b)** Three clusters (silhouette score = 0.87) encompass additives enriching pathways with cancer keywords, based on ERs for those pathways only. **c-e)** Clusters 1, 2, and 3 from **a**, respectively. Additives are sorted by IARC classification and display diverse usage patterns based on associations with the top polymer (PVC), function (colorant), and product category (food products). **f-h)** Clusters 1, 2, and 3 from **a** respectively. Each heatmap includes pathways from Wikipathways Cancer that are differentially enriched across the cluster. Pathways are divided by behavior and hierarchically clustered. Additives are hierarchically clustered by scaled ER and form subgroupings based on predicted cancer effects. **i)** Sankey diagrams characterize each cluster from **a**, highlighting their uniquely and highly enriched gene sets. Pie charts show that Clusters 1, 2, and 3 have near-identical distributions of carcinogenic, Group 3, and unclassified additives (although Cluster 3 lacks any Type 1 carcinogens).

To demonstrate how deeper connections between additives and cancer can be extracted from our dataset, we sorted the additives within each cluster into subgroups with similar cancer effects.

This was done through hierarchical clustering on the additives’ ERs for Wikipathways Cancer gene sets (**Fig. 4f-h**). Each *k*-means cluster exhibits a unique profile. Much of Cluster 1 (**Fig. 4f**) leans towards pathways impacting cell death/survival and DNA damage, with subgroups strongly impacting metabolic pathway WP143 (fatty acid beta-oxidation), immune pathway WP530 (cytokines and inflammatory response), and several cancer type pathways, specifically WP3859 (TGF-beta signaling in thyroid cells for epithelial-mesenchymal transition). Cluster 2 (**Fig. 4g**) appears to be broken into three main segments: the first subgroup strongly enriching cell cycle/proliferation pathway WP4357 (NRF2-ARE regulation) and DNA damage pathway WP3 (Transcriptional activation by NRF2 in response to phytochemicals); the second subgroup strongly enriching cell death/survival pathway WP3617 (Photodynamic therapy-induced NF-kB survival signaling) and slightly enriching cancer type pathway WP3859; and the third subgroup enriching cell death/survival pathway WP3617, immune pathway WP530, and cancer type pathway WP4337 (ncRNAs involved in STAT3 signaling in hepatocellular carcinoma). Cluster 3 (**Fig. 4h**) enriches cell death/survival, DNA damage, and cancer type pathways nearly across the board, with subgroups displaying particular enrichment for cell cycle/proliferation pathway WP4357, cell death/survival pathways WP3672 (lncRNA-mediated mechanisms of therapeutic resistance) and WP3617, DNA damage pathways WP1742 (TP53 network) and WP3, and immune pathway WP530. Even after hierarchical clustering by cancer-related effects, carcinogens and unclassified additives are interspersed (**Fig. 4f-h**).

Together, the affected pathways cover vast territory, including DNA damage, apoptosis, immune response, viral diseases, and cancer. Many pathways are affected by chemicals in all three clusters. However, distinguishing features of each individual cluster can be found through their unique and/or highly enriched pathways (**Fig. 4i**).

## Discussion

The pervasive nature of plastic and our frequent exposure to plastics has prompted increased attention to the potential harmful impacts of plastic on organismal health ^9^; however, many studies have focused on the influence of plastic polymers ^40^ or particularly well-studied additives, such as bisphenol A ^41^. Far less is known about the comprehensive landscape of plastic additives and mixtures of additives, including their environmental fates, transport, and their consequences for health and wellbeing. What little we know about additives is from studies on the additives in isolation, but these additives exist as complex mixtures of tens to hundreds of additives in a single plastic product (**Fig. 3e, 3g, Table S2, S7**), many of which exert widespread effects on gene expression (**Fig. 4f-i**). Here, we comprehensively characterize plastic additives for their potential carcinogenicity and impacts on gene expression. A striking observation from these analyses is the severe shortage of data on carcinogenic potential for hundreds of plastic additives (**Fig. 2a-c, 2e**). The apparent lack of documentation and cross-verification among databases raises questions about the efficacy of current legislation and safety measures for plastic and plastic additives (**Fig. 2d**).

Prior investigations have suggested that plastic ingestion may induce carcinogenesis. Studies in fish have shown that ingestion of microplastics induces hepatic inflammation ^42^ and hepatic neoplasia ^43^. Plastic contains multiple known carcinogens, the most well-studied of which is bisphenol A ^44^. At the cellular level, plastic exposure impacts numerous gene expression pathways linked to inflammatory signaling and cancer, including NF-kB ^45^, IL-6 ^46^, TNF alpha ^46^, and IL-8 ^47^. Consistent with these observations, our analyses revealed numerous impacts of plastic additives on gene expression pathways, many of which are relevant to cancer, including pro-inflammatory signaling and oxidative stress pathways (**Fig. 4f-i, Table S1, S4**). These effects on gene expression were exerted by both known carcinogens and additives for which carcinogenic potential is unknown, and clustering by gene expression pathways revealed substantial overlap between carcinogenic additives and additives with unknown carcinogenic potential (**Fig. 4)**. Further, clusters of additives enriched many converging phenotypes, such as proliferation and anti-apoptotic pathways, which may indicate greater carcinogenic risk (**Fig.4h**).

Our analyses provide a platform to pinpoint plastic products that harbor mixtures of additives with known consequences on gene expression that may impact human health. The substantial clustering of gene expression pathways (**Fig. 4a-b**) produced by carcinogenic and unclassified additives suggests that unclassified additives may be carcinogenic, though further testing and verification is required. These analyses may allow researchers and policymakers to identify and prioritize the populations and products that contain mixtures of additives with the greatest potential for harm. For example, we found 24 additives (including six known carcinogens and 12 unclassified additives) associated with plastics in construction materials that activated colorectal and gastric cancer pathways. Consistent with this, construction workers are at an enhanced risk for multiple cancer types, including esophageal, colorectal, gastric, and testicular cancer ^48^.

Whether these risks are associated with plastic additives requires further prospective interventional studies; however, our analysis provides a framework for identifying potential susceptible populations and associated products for follow-up study.

Our analyses have several limitations. There is an overall lack of transparency in the industrial literature regarding the presence of additives in common plastic polymers. Over 4,000 chemicals are estimated to be used in plastic food packaging, but our literature review only documented 2,712 additives – many of which lacked polymer and product data – across all applications ^28,49^. The key limitations in synthesizing exposure data were (1) misspelled, miscategorized, and unclear terms in the original review papers and (2) ambiguity when creating hierarchies of terms (*e*.*g*., “food-contact plastics”). Spelling corrections and grouping were either performed or checked manually because programmatic strategies like regex strings were prone to errors in prior research ^14^. However, manual database curation will not be scalable as research on plastic increases. A standardized and transparent way of disclosing, tracking, and reporting additives’ functions, polymers, and products will be necessary for the longevity of a comprehensive database.

Our toxicogenomics analysis revealed the presence of multiple carcinogenic additives in numerous plastic products. Perhaps more striking, however, is the severe lack of information about carcinogenic potential for the overwhelming majority of plastic additives. At the gene expression level, these unclassified additives impact many of the same pathways as known carcinogens. Collectively, these data underscore the critical need for a systematic study of plastic additives, with a focus on additives that overlap in their gene expression patterns with known carcinogens. We propose a transdisciplinary approach in which researchers, legislators, and manufacturers collaborate to address the following key gaps in our knowledge: 1) developing comprehensive toxicological profiles for individual plastic additives and common additive mixtures in plastic products, 2) mapping all additives to their functions and endpoints, 3) determining the fate and transport for individual additives and mixtures of additives that are leached from the same products in standardized settings, 4) determining toxicological synergies between groups of additives, and 5) identifying high-priority additives that should be removed or replaced to preserve plastic functionality. Our analyses seek to narrow the search space for carcinogenic additive combinations, which may accelerate reformulation strategies. Our work in documenting plastic additives can help guide future steps to enable a world where the health risks of plastics are both publicly known and effectively reduced.

## Methods

### Analytical workflow

We developed an analytical workflow consisting of the following steps: (1) literature-based review and identification of plastic additives, (2) characterization of additive coverage in public cancer databases, (3) integrated analysis of gene expression and exposure data, and cross-group comparisons. An *additive* is defined herein as any substance known to be added during the manufacturing process and/or detectable in the final polymer. Unexpected additives in the final polymer may be non-intentionally added substances during manufacturing or substances that adsorbed from the environment during and after use. Any chemical in a polymer could theoretically leach out and cause harmful health effects. Therefore, we deem it critical to include any chemical to which a human might be exposed when ingesting or contacting plastic. A parallel analysis was also conducted on polymers (*e*.*g*., polyethylene, polystyrene, polyurethane) to compare the extent of knowledge on additives vs. polymers. **Figure 1** provides an outline of the bioinformatics workflow for the project.

### Literature-based review and identification of plastic additives

To assemble a comprehensive list of plastic additives, we performed a literature review in Google Scholar, Clarivate Web of Science, and PubMed. In each database, targeted search strings (**Table S9**) were used to select peer-reviewed review articles containing lists of plastic additives (**Table S10**). Additives from each article were collected by their CAS numbers, assuming a one-to-one mapping of CAS number to substance. Several articles provided measures of confidence regarding the usage/presence of an additive; in these cases, only high-confidence additives were extracted. After seven articles were searched, the unique additive contribution per article began to diminish, plateauing at zero by the fourteenth paper (**Fig. S1**). In total, 18 articles produced 2,712 unique additives.

Sixteen papers compiled from the literature review included information about the function or purpose as a plastic additive (*e*.*g*., plasticizer, flame retardant), polymer usage (*e*.*g*., PET, PVA, PVC), and/or product usage (industry or consumer product, *e*.*g*., construction material, electronics, toys, textiles) of each additive. These data were manually collected in Excel and compiled using the pandas Python package (**Table S1**) ^50,51^. Ambiguous terms were included in all potential categories (*e*.*g*., when “adhesive” was found in a combined column of functions and products, it was recorded in both the Function and Product columns of our database). All function, polymer, and product strings can be found in **Table S11**.

Polymer names and acronyms were collected from the 18 literature review papers, other peer reviewed publications, and public listings of common polymers. They were mapped to CAS identifiers using PubChem. Alternate CAS numbers (if applicable) were retained and stored in a separate list from the primary number. There was no direct 1:1 mapping between any of the following variables: full chemical name, polymer acronym, and CAS number **(Table S2)**. To conduct the gene expression analysis, all chemicals under the same CAS number were grouped together.

### Characterization of additive coverage in public cancer databases

To evaluate the extent of accessible documentation on plastic additive carcinogenicity, three publicly available databases were selected: IRIS (Integrated Risk Information System from the U.S. EPA), IARC (International Agency for Research on Cancer), and ECOTOX (Ecotoxicology database from the U.S. EPA). Databases were queried using R Statistical Software (v4.2.1)^52^ and the tidyverse package ^53^. The IRIS database contains 651 chemicals, some of which are duplicated and have different carcinogenicity classifications depending on exposure route. If a single chemical was listed as carcinogenic and non-carcinogenic for different exposure routes, it was listed as carcinogenic for the analysis. As of September 2022, the IARC Database contained 1,101 total chemicals: 161 in Group 1 (carcinogenic to humans), 107 in Group 2A (probably carcinogenic to humans), 327 in Group 2B (possibly carcinogenic to humans), and 506 in Group 3 (inadequate evidence for carcinogenicity in humans). The IARC database was selected as the standard for categorizing chemicals prior to further downstream bioinformatics analyses. We considered chemicals in Groups 1, 2A, and 2B *carcinogens* in our analysis. The grouping of *carcinogens* and Group 3 chemicals were referred to as *classified* because all of these chemicals are classified in IARC and have had their carcinogenic potential evaluated. Any chemical lacking an IARC category is considered *unclassified* and has not been annotated with respect to its carcinogenic potential in IARC.

### Integrated analysis of gene expression and exposure data

Gene expression and pathway enrichment data were collected from Comparative Toxicogenomics Database (CTD), Mouse Genome Informatics (MGI), HUGO Gene Nomenclature Committee (HGNC) Comparison of Orthology Predictions (HCOP), and WebGestalt (WGA).

CTD is a public database sponsored by the National Institute of Environmental Health Sciences (NIEHS) which stores 50,048,577 toxicogenomic relationships. In this study, CTD was used to compile lists of human genes up-and down-regulated by each chemical additive. The database provided relationships for 289 unclassified chemicals and 139 classified chemicals. When polymer CAS numbers were separately screened through CTD, 29 substances (15 unique CAS numbers) were found to up-or down-regulate human genes according to published studies. On occasion, CTD labeled the interacting organism as human but erroneously provided a non-human GeneID. In these instances, the Mouse Genome Informatics Vertebrate Homology and HUGO Gene Nomenclature Committee Comparison of Orthology Predictions (HCOP) databases were queried to identify the corresponding human Entrez ID. If multiple human Entrez IDs were associated with one non-human homolog from CTD, all human matches were substituted for the homolog.

Using the gene lists from CTD, overrepresentation analysis (ORA) was performed in WebGestalt to predict pathway interactions for each additive (and polymer). For each substance, the ORA input was a single list combining all up-and down-regulated genes. The WebGestaltR library was used to conduct batch ORA for all 428 additives and 15 polymers (polymers under the same CAS were grouped together). The PANTHER, Reactome, KEGG, Wikipathway, and Wikipathway Cancer pathway databases were queried with an FDR threshold of 25% and a maximum of 2,000 genes (default). Results were generated for seven polymers (46.67%), 120 classified chemicals (86.33%, 65% of which were carcinogens), and 229 unclassified chemicals (79.24%) (**Tables S1, S2**). Even after homology correction, a small percentage of genes from CTD (<1%) remained unmappable in WebGestalt, but the majority of gene identifiers were recognized.

### Clustering and cross-group comparisons

Dimensionality reduction and clustering were performed for each plastic additive in the Python programming language using sklearn.cluster.KMeans, sklearn.decomposition.PCA, and umap.umap_. A pairwise matrix of Enrichment Ratio (ER) for each plastic additive and pathway was constructed to facilitate weighted clustering on pathway enrichment. Principal Component Analysis (PCA) was performed to achieve 95% explained variance with minimal dimensionality. The resulting matrix was further reduced using Uniform Manifold Approximation and Projection (UMAP) to improve clustering results.

Cluster quality was heavily dependent on the two nondeterministic algorithms in this workflow: UMAP and *k*-means. Running *k*-means after applying default UMAP parameters (n_neighbors = 40, n_components = 2, min_dist = 0.3) was not sufficient for any *k*, producing silhouette scores with low and sometimes negative values. Silhouette scores below zero indicate that elements have been assigned to the wrong clusters; scores near zero indicate that clusters overlap; scores near 1 indicate that most elements cluster more closely within their assigned cluster than other clusters. To ensure high-quality clusters, a minimum silhouette score of 0.70 was selected.

UMAP’s n_neighbors parameter was tested at all integer values between 2 and 20 inclusive, while min_dist was tested at values 0.0, 0.1, 0.25, 0.5, 0.8, and 0.99. We carried out *k*-means for all *k* between 3 and 10, and the number of clusters producing the optimal silhouette score was selected. The random states for both UMAP and *k*-means were modulated between five different values to capture a broader range of possible results. This procedure was repeated for (1) all additives and all ERs, (2) only additives enriching at least one pathway with a cancer keyword substring [‘cancer’, ‘carcin’, ‘metasta’, ‘tumor’] and the ERs for those pathways, and (3) only additives enriching at least one pathway with an oxidative stress keyword [‘oxidative stress’, ‘oxidative damage’, ‘reactive oxygen species’, ‘ros’] and the ERs for those pathways (**Table S1**). In all three cases, the vast majority of top-performing cluster runs produced three to four clusters (**Table S6**). The consistent clustering results for much lower-dimensionality clustering (cancer keyword and oxidative-stress keyword pathways only) validated the three-cluster result for all additives and all ERs.

All subsequent analyses were performed on the three clusters made from the full dataset. To determine subgroupings with similar cancer effects, Wikipathways Cancer pathways differentially enriched across a cluster above a certain standard deviation threshold (10, 10, 20 for clusters 1, 2, and 3 respectively) were selected. Only additives enriching at least one of those pathways were retained. Enrichment ratios were scaled using the scale() function in R, and both additives and pathways were hierarchically clustered using ComplexHeatmap. Each pathway was manually assigned to one cancer-relation category (cancer type, cell cycle/proliferation, cell death/survival, DNA damage, immune, metabolism) based on its most prominent effects according to Wikipathways and published literature.

To distinguish the most salient pathways for each cluster, a binary matrix was constructed to indicate whether each additive in the cluster enriched or did not enrich a particular pathway. The 50 pathways with the most additive associations were considered the central pathways for that cluster. “Uniquely enriched” pathways for a cluster do not appear in any other cluster’s top 50. “Highly enriched” pathways for a cluster have at least one ER >=100.

## Data and Code Availability

All code will be made available upon request. See Supplementary File 1 and 2 for all data.

## Acknowledgements

We would like to thank Bass Connections at Duke University for funding this research. We would also like to thank the Duke Plastic Pollution Working Group, the Duke Nicholas Institute for Energy, Environment, & Sustainability, and the Duke University Marine Lab Scholars.

## Author Contributions

JAS supervised the study. ZD and JAS conceived the study. SV, JS, BS, and MM collected additive and polymer CAS numbers and finalized the study design. BS identified and queried chemicals through three chemical databases. SV collected exposure and gene expression data, performed enrichment analysis and clustering, and assembled supplementary tables. JS analyzed and compared additive clusters. SV, BS, and JS wrote the first draft of the manuscript. All authors collaborated to write, edit, and approve the final version of the manuscript.

## Additional Information

Additional information can be found within the Supplementary Materials.

Reprints and permissions information is available at www.nature.com/reprints.

## References

1. Thompson RC, Swan SH, Moore CJ, vom Saal FS. Our plastic age. Philosophical Transactions of the Royal Society B: Biological Sciences. 2009;364(1526):1973–1976. doi:10.1098/rstb.2009.0054

2. Geyer R, Jambeck JR, Law KL. Production, use, and fate of all plastics ever made. Science Advances. 2017;3(7):e1700782. doi:10.1126/sciadv.1700782

3. Ellen MacArthur Foundation. The New Plastics Economy: Rethinking the future of plastics. Published 2016.Accessed October 12, 2022. https://ellenmacarthurfoundation.org/the-new-plastics-economy-rethinking-the-future-of-plastics

4. Grand View Research. Plastic Market Size, Share & Trends Report, 2022 - 2030.; 2019:230.Accessed March 17, 2023. https://www.grandviewresearch.com/industry-analysis/global-plastics-market

5. PlasticsEurope. Plastics - the Facts 2019. Published online 2019.Accessed October 12, 2022. https://plasticseurope.org/wp-content/uploads/2021/10/2019-Plastics-the-facts.pdf

6. Andrady AL, Neal MA. Applications and societal benefits of plastics. Philosophical Transactions of the Royal Society B: Biological Sciences. 2009;364(1526):1977–1984. doi:10.1098/rstb.2008.0304

7. Barboza LGA, Dick Vethaak A, Lavorante BRBO, Lundebye AK, Guilhermino L. Marine microplastic debris: An emerging issue for food security, food safety and human health. Marine Pollution Bulletin. 2018;133:336–348. doi:10.1016/j.marpolbul.2018.05.047

8. DeWeerdt S. How to make plastic less of an environmental burden. Nature. 2022;611(7936):S2–S5. doi:10.1038/d41586-022-03644-1

9. Morrison M, Trevisan R, Ranasinghe P, et al. A growing crisis for One Health: Impacts of plastic pollution across layers of biological function. Frontiers in Marine Science. 2022;9.Accessed March 17, 2023. https://www.frontiersin.org/articles/10.3389/fmars.2022.980705

10. Rodrigues MO, Abrantes N, Gonçalves FJM, Nogueira H, Marques JC, Gonçalves AMM. Impacts of plastic products used in daily life on the environment and human health: What is known? Environmental Toxicology and Pharmacology. 2019;72:103239. doi:10.1016/j.etap.2019.103239

11. Thompson T. Plastic pollution: Three problems that a global treaty could solve. Nature. Published online November 28, 2022. doi:10.1038/d41586-022-03835-w

12. Vethaak AD, Leslie HA. Plastic Debris Is a Human Health Issue. Environ Sci Technol. 2016;50(13):6825–6826. doi:10.1021/acs.est.6b02569

13. Wright SL, Kelly FJ. Plastic and Human Health: A Micro Issue? Environ Sci Technol. 2017;51(12):6634–6647. doi:10.1021/acs.est.7b00423

14. Wiesinger H, Wang Z, Hellweg S. Deep Dive into Plastic Monomers, Additives, and Processing Aids. Environ Sci Technol. 2021;55(13):9339–9351. doi:10.1021/acs.est.1c00976

15. Habibi N, Uddin S, Fowler SW, Behbehani M. Microplastics in the atmosphere: a review. Journal of Environmental Exposure Assessment. 2022;1(1):6. doi:10.20517/jeea.2021.07

16. Jahnke A, Arp HPH, Escher BI, et al. Reducing Uncertainty and Confronting Ignorance about the Possible Impacts of Weathering Plastic in the Marine Environment. Environ Sci Technol Lett. 2017;4(3):85–90. doi:10.1021/acs.estlett.7b00008

17. Sobhani Z, Lei Y, Tang Y, et al. Microplastics generated when opening plastic packaging. Sci Rep. 2020;10(1):4841. doi:10.1038/s41598-020-61146-4

18. Baj J, Dring JC, Czeczelewski M, et al. Derivatives of Plastics as Potential Carcinogenic Factors: The Current State of Knowledge. Cancers. 2022;14(19):4637. doi:10.3390/cancers14194637

19. Cox KD, Covernton GA, Davies HL, Dower JF, Juanes F, Dudas SE. Human Consumption of Microplastics. Environ Sci Technol. 2019;53(12):7068–7074. doi:10.1021/acs.est.9b01517

20. Lett Z, Hall A, Skidmore S, Alves NJ. Environmental microplastic and nanoplastic: Exposure routes and effects on coagulation and the cardiovascular system. Environmental Pollution. 2021;291:118190. doi:10.1016/j.envpol.2021.118190

21. Amato-Lourenco LF, Carvalho-Oliveira R, Ribeiro Junior G, Galvao L dos S, Ando RA, Mauad T. Presence of airborne microplastics in human lung tissue. J Hazard Mater. 2021;416:126124. doi:10.1016/j.jhazmat.2021.126124

22. Ragusa A, Svelato A, Santacroce C, et al. Plasticenta: First evidence of microplastics in human placenta. Environ Int. 2021;146:106274. doi:10.1016/j.envint.2020.106274

23. Ragusa A, Notarstefano V, Svelato A, et al. Raman Microspectroscopy Detection and Characterisation of Microplastics in Human Breastmilk. Polymers (Basel). 2022;14(13):2700. doi:10.3390/polym14132700

24. Yusof SI, Anuar ST, Azmi AA, et al. Detection of microplastics in human colectomy specimens. JGH Open. 2021;5(1):116–121. doi:http://dx.doi.org/10.1002/jgh3.12457

25. Yates J, Deeney M, Rolker HB, White H, Kalamatianou S, Kadiyala S. A systematic scoping review of environmental, food security and health impacts of food system plastics. Nat Food. 2021;2(2):80–87. doi:10.1038/s43016-021-00221-z

26. Rochman CM, Brookson C, Bikker J, et al. Rethinking microplastics as a diverse contaminant suite. Environmental Toxicology and Chemistry. 2019;38(4):703–711. doi:https://doi.org/10.1002/etc.4371

27. Sendra M, Pereiro P, Figueras A, Novoa B. An integrative toxicogenomic analysis of plastic additives. Journal of Hazardous Materials. 2021;409:124975. doi:10.1016/j.jhazmat.2020.124975

28. Groh KJ, Backhaus T, Carney-Almroth B, et al. Overview of known plastic packaging-associated chemicals and their hazards. Science of The Total Environment. 2019;651:3253–3268. doi:10.1016/j.scitotenv.2018.10.015

29. Campanale C, Massarelli C, Savino I, Locaputo V, Uricchio VF. A Detailed Review Study on Potential Effects of Microplastics and Additives of Concern on Human Health. Int J Environ Res Public Health. 2020;17(4):1212. doi:10.3390/ijerph17041212

30. Hirai H, Takada H, Ogata Y, et al. Organic micropollutants in marine plastics debris from the open ocean and remote and urban beaches. Marine Pollution Bulletin. 2011;62(8):1683–1692. doi:10.1016/j.marpolbul.2011.06.004

31. Hahladakis JN, Velis CA, Weber R, Iacovidou E, Purnell P. An overview of chemical additives present in plastics: Migration, release, fate and environmental impact during their use, disposal and recycling. Journal of Hazardous Materials. 2018;344:179–199. doi:10.1016/j.jhazmat.2017.10.014

32. Hartmann NB, Hüffer T, Thompson RC, et al. Are We Speaking the Same Language? Recommendations for a Definition and Categorization Framework for Plastic Debris. Environ Sci Technol. 2019;53(3):1039–1047. doi:10.1021/acs.est.8b05297

33. Jang M, Shim WJ, Han GM, Rani M, Song YK, Hong SH. Styrofoam Debris as a Source of Hazardous Additives for Marine Organisms. Environ Sci Technol. 2016;50(10):4951–4960. doi:10.1021/acs.est.5b05485

34. Barboza LGA, Cunha SC, Monteiro C, Fernandes JO, Guilhermino L. Bisphenol A and its analogs in muscle and liver of fish from the North East Atlantic Ocean in relation to microplastic contamination. Exposure and risk to human consumers. Journal of Hazardous Materials. 2020;393:122419. doi:10.1016/j.jhazmat.2020.122419

35. Rochman CM, Lewison RL, Eriksen M, Allen H, Cook AM, Teh SJ. Polybrominated diphenyl ethers (PBDEs) in fish tissue may be an indicator of plastic contamination in marine habitats. Science of The Total Environment. 2014;476–477:622-633. doi:10.1016/j.scitotenv.2014.01.058

36. Tanaka K, Takada H, Yamashita R, Mizukawa K, Fukuwaka M aki, Watanuki Y. Accumulation of plastic-derived chemicals in tissues of seabirds ingesting marine plastics. Marine Pollution Bulletin. 2013;69(1):219–222. doi:10.1016/j.marpolbul.2012.12.010

37. Fossi MC, Marsili L, Baini M, et al. Fin whales and microplastics: The Mediterranean Sea and the Sea of Cortez scenarios. Environmental Pollution. 2016;209:68–78. doi:10.1016/j.envpol.2015.11.022

38. Hu X, Biswas A, Sharma A, et al. Mutational signatures associated with exposure to carcinogenic microplastic compounds bisphenol A and styrene oxide. NAR Cancer. 2021;3(1):zcab004. doi:10.1093/narcan/zcab004

39. Kim H, Zaheer J, Choi EJ, Kim JS. Enhanced ASGR2 by microplastic exposure leads to resistance to therapy in gastric cancer. Theranostics. 2022;12(7):3217–3236. doi:10.7150/thno.73226

40. Lithner D, Larsson Å Dave G. Environmental and health hazard ranking and assessment of plastic polymers based on chemical composition. Science of The Total Environment. 2011;409(18):3309–3324. doi:10.1016/j.scitotenv.2011.04.038

41. Jalal N, Surendranath AR, Pathak JL, Yu S, Chung CY. Bisphenol A (BPA) the mighty and the mutagenic. Toxicol Rep. 2017;5:76–84. doi:10.1016/j.toxrep.2017.12.013

42. Zhao L, Shi W, Hu F, Song X, Cheng Z, Zhou J. Prolonged oral ingestion of microplastics induced inflammation in the liver tissues of C57BL/6J mice through polarization of macrophages and increased infiltration of natural killer cells. Ecotoxicology and Environmental Safety. 2021;227:112882. doi:10.1016/j.ecoenv.2021.112882

43. Rochman CM, Hoh E, Kurobe T, Teh SJ. Ingested plastic transfers hazardous chemicals to fish and induces hepatic stress. Sci Rep. 2013;3:3263. doi:10.1038/srep03263

44. Murray TJ, Maffini MV, Ucci AA, Sonnenschein C, Soto AM. Induction of mammary gland ductal hyperplasias and carcinoma in situ following fetal bisphenol A exposure. Reprod Toxicol. 2007;23(3):383–390. doi:10.1016/j.reprotox.2006.10.002

45. Zhang Y, Yin K, Wang D, et al. Polystyrene microplastics-induced cardiotoxicity in chickens via the ROS-driven NF-κB-NLRP3-GSDMD and AMPK-PGC-1α axes. Sci Total Environ. 2022;840:156727. doi:10.1016/j.scitotenv.2022.156727

46. Hwang J, Choi D, Han S, Choi J, Hong J. An assessment of the toxicity of polypropylene microplastics in human derived cells. Sci Total Environ. 2019;684:657–669. doi:10.1016/j.scitotenv.2019.05.071

47. Weber A, Schwiebs A, Solhaug H, et al. Nanoplastics affect the inflammatory cytokine release by primary human monocytes and dendritic cells. Environment International. 2022;163:107173. doi:10.1016/j.envint.2022.107173

48. Fucic A, Galea KS, Duca RC, et al. Potential Health Risk of Endocrine Disruptors in Construction Sector and Plastics Industry: A New Paradigm in Occupational Health. International Journal of Environmental Research and Public Health. 2018;15(6). doi:10.3390/ijerph15061229

49. Muncke J, Andersson AM, Backhaus T, et al. Impacts of food contact chemicals on human health: a consensus statement. Environmental Health. 2020;19(1):25. doi:10.1186/s12940-020-0572-5

## Methods References

50. McKinney W. Data Structures for Statistical Computing in Python. Proceedings of the 9th Python in Science Conference. Published online 2010:56-61. doi:10.25080/Majora-92bf1922-00a

51. The pandas development team. pandas-dev/pandas: Pandas. https://doi.org/10.5281/zenodo.3509134

52. R Core Team. R: A language and environment for statistical computing. Published online 2022. https://www.R-project.org/

53. Wickham H, Averick M, Bryan J, et al. Welcome to the tidyverse. Journal of Open Source Software. 2019;4(43):1686. doi:10.21105/joss.01686

