## Supplemental Materials for "Toxicogenomic analysis of the carcinogenic potential of plastic additives"

### Supplementary Material

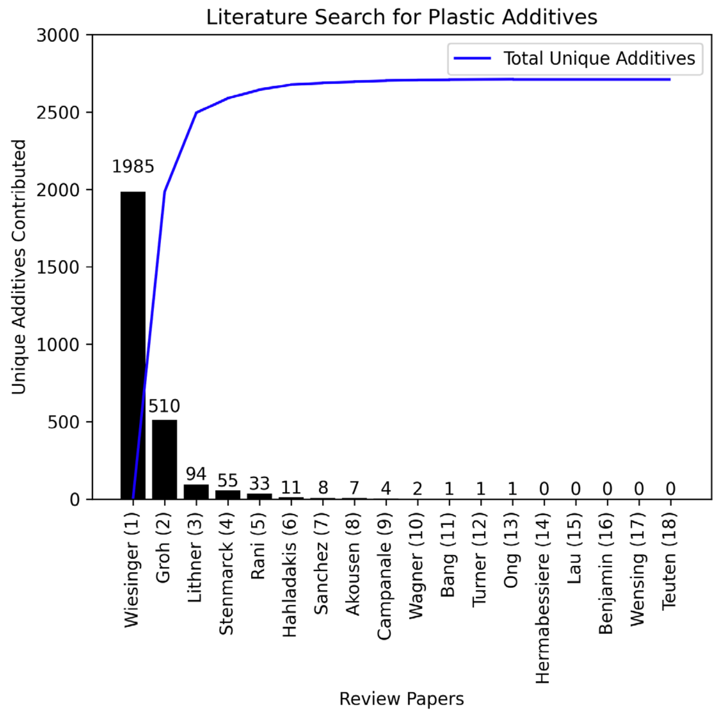


**Figure S1:** Literature search for plastic additives. The search was limited to 18 review papers as the number of unique additives discovered reached a consistent plateau.

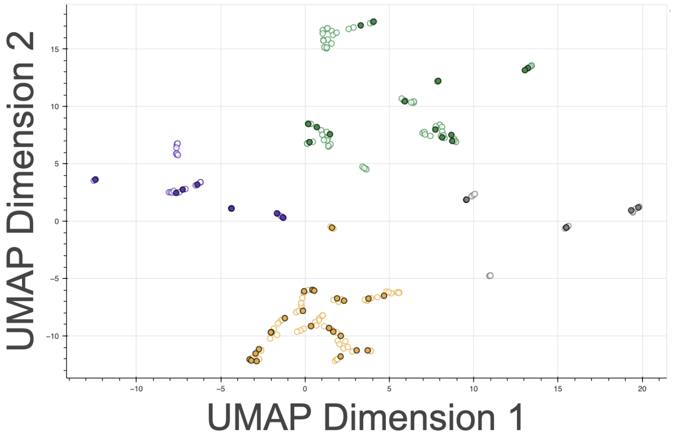


**Figure S2:** Four clusters form when only enrichment scores for pathways with oxidative stress keywords [‘OXIDATIVE STRESS’, ‘OXIDATIVE DAMAGE’, ‘REACTIVE OXYGEN SPECIES’, ‘ROS’]. Only seven pathways were used: ['OXIDATIVE DAMAGE_W', 'OXIDATIVE STRESS INDUCED SENESCENCE_R', 'OXIDATIVE STRESS RESPONSE_P', 'OXIDATIVE STRESS_W', 'DETOXIFICATION OF REACTIVE OXYGEN SPECIES_R', 'SIMPLIFIED INTERACTION MAP BETWEEN LOXL4 AND OXIDATIVE STRESS PATHWAY_W', 'ETHANOL METABOLISM RESULTING IN PRODUCTION OF ROS BY CYP2E1_W']

**Supplementary File Key**

**Supplementary File 1** contains Table S1 and S2. **Supplementary File 2** contains Tables S3-S11.

**Table S1.** All information collected on plastic additives. Columns = 'CAS', 'Function', 'Polymers', 'Products', 'Reprocessed' (indicating whether a CAS number was reformatted to standard formatting, *e.g.,* 10-2-2 → 10-02-2), 'Source', 'Name', 'In CTD', 'Polymers CAS', 'Total Enriched Pathways', 'Enriched Pathways', 'Upregulated Genes (Symbol)', 'Upregulated Genes (Entrez ID)', 'Total Upregulated Genes', 'Downregulated Genes (Symbol)', 'Downregulated Genes (Entrez ID)', 'Total Downregulated Genes', 'PubMedIDs on Gene Interactions', 'Total Publications on Gene Interactions', 'Known Carcinogenicity', 'IARC Category', 'In IRIS', 'IRIS_assessment_type', 'IRIS_WOE_characterization', 'In Ecotox', 'ecotox_carcinogenic_effect_and_measurement', 'Total Polymer Types', 'Total Functions', 'Cluster All ERs', 'Total Products', 'Total Polymers CAS'

**Table S2.** Full polymer data. Columns = 'Acronym', 'Name', 'Reference Name', 'Primary CAS', 'All CAS', 'In CTD', 'Upregulated Genes (Symbol)', 'Upregulated Genes (Entrez ID)', 'Total Upregulated Genes', 'Downregulated Genes (Symbol)', 'Downregulated Genes (Entrez ID)', 'Total Downregulated Genes', 'PubMedIDs', 'Publications', 'Enriched Pathways', 'Enriched Pathway IDs', 'Total Enriched Pathways', 'Additives', 'Total Additives',

**Table S3.** Product categorizations. Columns = 'Product Category', 'Additives', 'Search Strings', 'Products'

**Table S4.** Top ten upregulated genes, downregulated genes, and enriched gene sets for carcinogens vs. unclassified additives.

**Table S5.** Student t-tests comparing data availability between additives in different IARC categories. Significance is noted where $p < 0.05$.

**Table S6.** Silhouette scores provide evidence for $k=3$ clusters in $k$-means.

**Table S7.** Full product data, containing 415 unique product strings. Columns = 'Product', 'Additives', 'Total Additives'

**Table S8.** Full function data, containing 168 unique function strings. Columns = 'Function', 'Additives', 'Total Additives'

**Table S9.** Search strings used to locate review articles that provide chemical additives intentionally used in and/or consistently found in plastics.

**Table S10.** Review papers used to curate lists of plastic additives and exposure data for each additive.

**Table S11.** All categorizations. Columns = Function, Polymers, Products. There are 279 unique polymer strings, 168 unique function strings, and 415 unique product strings.
